# Donor-Free Gene Correction by Targeted Interhomolog Recombination

**DOI:** 10.1101/538603

**Authors:** Luther Davis, Kevin J. Khoo, Nancy Maizels

## Abstract

Spontaneous gene correction by interhomolog recombination (IHR) occasionally occurs to ameliorate genetic diseases of blood and skin^1–3^. Using an engineered endogenous gene as a reporter, we demonstrate that gene correction by IHR is normally infrequent (≤0.02%) but is stimulated by DSBs targeted by CRISPR/Cas9 to both homologous chromosomes; reaching frequencies of 0.5%. We further show that depletion of POLQ stimulates IHR frequencies 4-fold, to 2%, and promotes IHR in G2 phase, when recombination between replicated homologs can correct not only compound heterozygous but also autosomal dominant “gain-of-function” mutations, which present a special challenge for gene therapy. The strategies reported here will enable optimization of IHR for gene therapy in a variety of cell types. Advantages include the ability to correct gain-of-function mutations, no need for an exogenous donor, and the potential to limit damage to coding sequence by targeting IHR to introns.

---

In order to systematically quantify and optimize IHR, we engineered the endogenous CD44 gene in human HT1080 cells to report on IHR events by flow cytometry. HT1080 cells provide an appropriate model as they express functional P53, maintain a diploid karyotype in culture^4, 5^, and their high cloning efficiency simplifies molecular analysis of recombinants. The CD44 gene, which contains 18 exons and spans 94 kb on chromosome 11p, was modified to derive a robust and readily generalizable reporter by targeted mutation of exon 1 on one allele and exon 17 on the other (**Fig. 1a, top**). CD44 encodes a cell surface glycoprotein expressed by most cells. Cells that express functional protein (sCD44+) can be scored by flow cytometry after staining with commercial antibody. Crossover recombination at sites between the mutations in exons 1 and 17 in the engineered HT1080-K1 CD44-/- line generates sCD44+ cells with two distinct genotypes (**Fig. 1a, below**). IHR resulting in reciprocal exchange of genetic information will generate one homolog with no mutations and another with two mutations (**left**). IHR resulting in copy-neutral loss of heterozygosity (LOH) will substitute correct sequence for the telomere proximal mutation in exon 1 in one daughter cell (**right**). This outcome of IHR, which can correct autosomal dominant (“gain of function”) mutations as illustrated in **Supplementary Fig. 1**, was quantified by RFLP analysis of sCD44+ recombinants.

**Figure 1.**
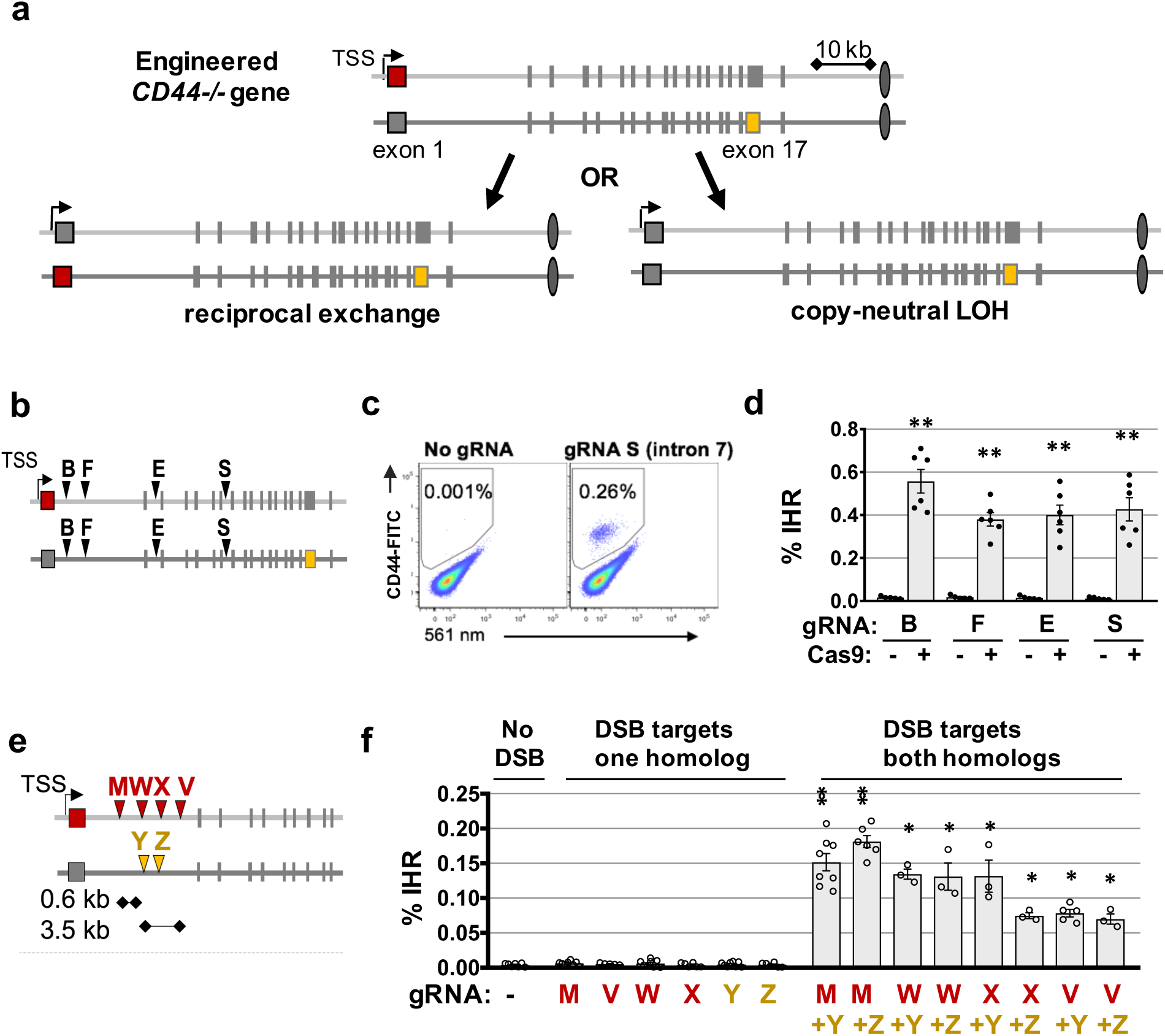
Correction of mutations by IHR at targeted DSBs. (a) Above, diagram of the two alleles of the engineered CD44 gene in CD44-/- HT1080 derivatives, mutant in exon 1 (red) or 17 (gold). Below: Homologs generated following IHR that results in reciprocal exchange (left) or copy-neutral LOH of sequences telomeric to the recombination site (right). (b) Map of naturally-occurring gRNA sites B, F, E and S (arrowheads) present on both alleles of the engineered CD44-/- gene in HT1080-K1 cells. (c) Representative flow cytometry analysis of HT1080-K1 populations targeted by Cas9 and no gRNA, or gRNA S that targets CD44 intron 7. Fraction of sCD44+ cells in boxed gates. (d) Frequencies of IHR in HT1080-K1 cells in which DSBs were targeted to intronic sites in both homologs by gRNAs B, F, E and S. Control cells transfected with expression constructs for gRNA but not Cas9. IHR frequencies are shown as mean and standard error of the mean (SEM, n≥5); ** indicate significant difference from the corresponding no Cas9 control (p<1×10-3). (e) Map of gRNA sites (arrowheads) present on only a single homolog (upper, red; lower, gold) in the engineered CD44-/- gene in HT1080-K2 cells. (f) Frequencies of IHR in HT1080-K2 cells in which IHR was initiated by DSBs targeted to a site on only one homolog; or to offset sites on both homologs. IHR frequencies are corrected for transfection efficiency and shown as mean and SEM (n≥3); * and ** indicate significant difference from no gRNA control (p<0.05 and p<10-5, respectively).

The frequency of sCD44+ cells detected by flow cytometry in normally proliferating cultures of engineered CD44-/- HT1080-K1 cells was <0.005%. We tested the ability of targeted DSBs to stimulate IHR by transfection with constructs expressing Cas9 and gRNAs B, F, E or S that direct cleavage to naturally-occurring intronic sites on both homologs of the CD44 gene (**Fig. 1b**). Targeted DSBs stimulated IHR to generate a sCD44+ cell population readily scored by flow cytometry (e.g. **Fig. 1c**). Correcting for transfection efficiency (30%), frequencies of IHR at targeted DSBs ranged from 0.38-0.56%, significantly above the background in control populations expressing only gRNA (p<4 x 10^-4^; **Fig. 1d**). Analysis of junctions from sCD44+ recombinants identified indels at 27/29, consistent with use of an end-joining pathway.

In order for IHR to proceed by reciprocal end-joining DSBs must be targeted to both homologs. In contrast, a DSB targeted to a single homolog will be sufficient to initiate homologous recombination. To distinguish these pathways, intron 1 of the CD44 gene was engineered to carry target sites unique to individual homologs, generating the HT1080-K2 derivative which carries cleavage sites M, W, X and V on one allele and Y and Z on the other (**Fig. 1e**). DSBs targeted to only a single homolog (by gRNAs M, V, W, X, Y or Z) did not stimulate IHR to reach frequencies significantly different than frequencies observed in the absence of gRNA (p>0.1; **Fig. 1f**). In contrast, DSBs targeted to different sites on the two homologs (“offset” rather than “aligned” sites) stimulated IHR to reach frequencies in the range of 0.23-0.60%, significantly above frequencies at DSBs targeted to a single homolog (p<0.03; **Fig. 1f**). Frequencies were comparable at offset DSBs separated by as little as 3 bp (gRNAs W+Y, W+Z, X+Y and X+Z) and by as much as 0.6 kb (by gRNA pairs M+Y or M+Z) or 3.5 kb (by gRNA pair V+Y or V+Z). IHR initiated by offset DSBs is predicted to generate clones in which the region flanked by the DSBs is deleted in one homolog and has undergone duplication in the other (**Supplementary Fig. 2**). PCR amplification of recombination junctions in individual sCD44+ recombinants resulting from IHR initiated by gRNAs M and Y, which cleave at sites offset by 0.6 kb, identified clones carrying these predicted deletions/tandem duplications, as well as clones in which the amplified region was the size of the parental DNA, as predicted for products of homologous recombination, in a ratio of approximately 5:1.

RFLP analysis showed that frequencies of LOH in exon 1 were similar among recombinants initiated by aligned and offset DSBs [5.8% (6 of 104) and 4.6% (4/88), respectively. RFLP analysis of SNPs rs85074 and rs7950932, which map 3.1 and 33.2 Mb telomeric of CD44, showed that there was complete correlation between homozygosity at exon 1 and homozygosity at these SNPs (rs85074, 5 of 5; rs7950932, 4 of 4), and heterozygosity at exon 1 and heterozygosity at these SNPs (rs85074, 67 of 67; rs7950932, 24 of 24). Exon 17, which is centromere proximal, was heterozygous in all 72 clones tested, including 5 that were homozygous at exon 1. These results demonstrate that a small fraction of IHR results in LOH, which extends from the site of recombination to the telomere.

The contribution of homologous recombination to IHR was further addressed by depletion of homologous recombination factors BRCA2, which promotes canonical HDR by loading RAD51 on resected DNA ends^6^; or RAD52, which supports strand annealing in pathways that are independent of BRCA2/RAD51. Treatment with either siBRCA2 or siRAD52 prior to transfection with Cas9 RNPs did not significantly affect IHR frequencies at either aligned or offset DSBs (**Fig. 2a**). These results, like the molecular analysis above, indicate that only a minor fraction of IHR events occur by homologous recombination.

**Figure 2.**
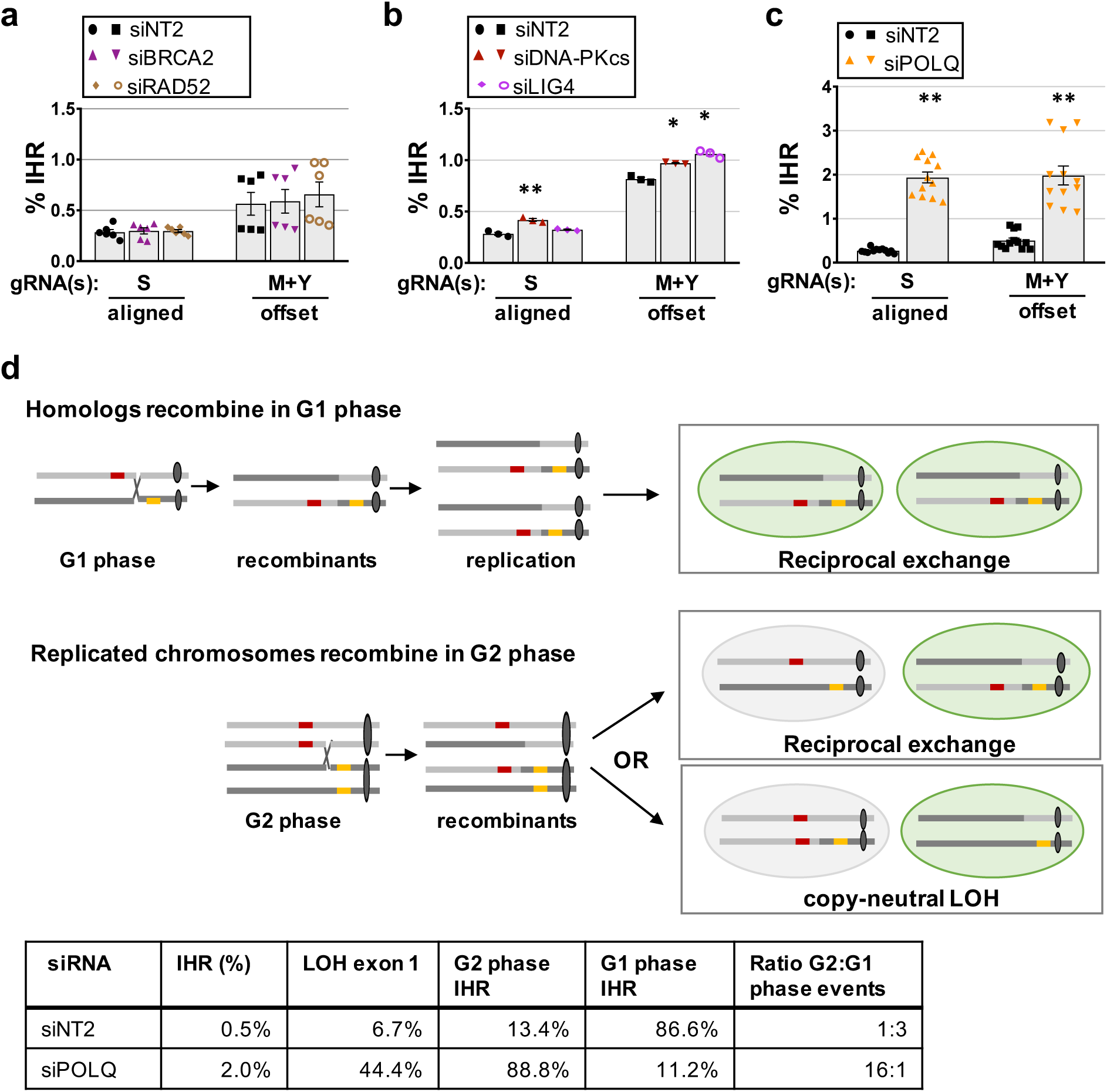
POLQ depletion stimulates G2 phase IHR. (**a-c**). Frequencies of IHR in HT1080-K2 cells treated with indicated siRNAs and transduced with RNPs composed of Cas9 and indicated gRNAs. siNT2, non-targeting control. IHR frequencies are shown as mean and SEM (n≥3); * and ** indicate significant difference from siNT2 (p<0.05 and p<10-3, respectively). (a) siNT2, siBRCA2, siRAD52 (b) siNT2, siDNA-PKcs, siLIG4 (c) siNT2, siPOLQ. (d) IHR outcomes in G1 and G2 phases. Above: IHR in G1 phase of cell cycle corrects a compound heterozygous mutation. Following segregation, the two identical daughter cells will each contain one homolog that has undergone reciprocal recombination and one in parental configuration, and both will express functional protein (green shading). IHR between replicated homologs in G2 phase can have two outcomes. If both recombinant chromosomes segregate to the same daughter cell, the compound heterozygous mutation will be corrected in that daughter (green shading) but not the other, which will retain the parental genotype. Alternatively, if the recombinant chromosomes segregate to different daughter cells, the compound heterozygous mutation will be corrected in one of those cells (green shading) and both daughter cells will exhibit copy-neutral LOH along the region that extends from the recombination junction to the telomere. LOH can correct gain-of-function mutants, as diagrammed in detail in **Supplementary Fig. 1**. Below: tabulation of the effect of POLQ depletion on cell cycle regulation of IHR at offset DSBs.

The contribution of canonical nonhomologous end-joining (c-NHEJ) was tested by depletion of the DNA-Protein Kinase catalytic subunit (DNA-PKcs) and DNA ligase 4 (LIG4)^7, 8^ (**Fig. 2b**). Depletion of DNA-PKcs caused a modest but significant increase in IHR frequencies at both offset and aligned DSBs (1.2- and 1.3-fold, respectively; p = 0.008 and 0.003). Depletion of LIG4 also caused a modest but significant increase in IHR frequency at offset (1.5-fold; p = 0.001) but not aligned (1.1-fold; p = 0.11) DSBs. Thus, c-NHEJ factors modestly inhibit IHR at targeted DSBs, perhaps by promoting rejoining of cleaved ends in cis.

The contribution of alternative end joining (altEJ) was tested by depletion of POLQ, which encodes DNA Polθ, a helicase and translesion polymerase that plays a critical role in repair of DSBs formed by a variety of mechanisms, including stalled replication forks, CRISPR/Cas9 cleavage and ionizing radiation^8–15^. Depletion of Polθ has been reported to inhibit altEJ but stimulate homologous recombination^16, 17^. Strikingly, depletion of POLQ stimulated IHR 4-fold or more, increasing IHR frequencies to 2% at both offset and aligned DSBs (**Fig. 2c**; p = 2.0 x 10^-5^ and 2.9 x 10^-8^). RFLP analysis scored LOH at exon 1 in 6.7% (5/75) of sCD44+ recombinants from siNT2-treated control cells, and in 44.4% (24/54) of sCD44+ recombinants from cells treated with siPOLQ (p<0.0001, two-tailed Fisher’s exact test). Thus, depletion of POLQ boosted IHR frequencies, especially IHR resulting in LOH.

IHR in G2 phase but not G1 phase can result in LOH (**Fig. 2d**). The fraction of recombinants that derive from G1 and G2 phase IHR events was estimated based on the frequency of LOH among sCD44+ recombinants. Assuming that recombinant chromosomes are equally likely to segregate to the same or different daughter cells following G2 phase IHR, then there will be equal numbers of sCD44+ recombinants produced in G2 phase that do and do not exhibit LOH (**Fig. 2d**). Based on this, we estimate that in cells treated with control siNT2, 13.4% (2 x 6.7%) of recombinants were generated in G2 phase, and the remaining 86.6% in G1 phase; and in cells depleted for POLQ, 88.8% (2 x 44.4%) of recombinants were generated in G2 phase and 11.2% in G1 phase (**Fig. 2e**).

The relative frequencies of G1 and G2 phase IHR events was estimated after dividing the fraction of G1 phase recombinants by two, as each G1 phase recombination event produces two daughters that are sCD44+, while each G2 phase event produces only one (**Fig. 2d**). The ratio of IHR events in G2:G1 phase was approximately 1:3 (13.4% relative to 86.6%/2=43.3%) in siNT2-treated control cells and 16:1 (88.8% relative to 11.2%/2 = 5.6%) in siPOLQ-treated cells. Thus, POLQ depletion caused G2 phase events to increase 48-fold (**Fig. 2e**). These results suggest that Polθ normally plays a critical role in ensuring that unrepaired DSBs do not persist into G2 phase, to limit IHR accompanied by LOH and also, perhaps, to limit translocation.

Identification of Polθ downregulation as a novel strategy to stimulate IHR between replicated homologs in G2 phase enhances the ability to correct autosomal dominant mutations, which account for a significant fraction of human genetic disease and have presented a particular challenge as these diseases are not candidates for gene replacement therapy. Accompanying copy-neutral LOH may limit the applicability of this strategy in some contexts, nonetheless the evidence that spontaneous gene correction ameliorates some diseases^1–3^ clearly indicates that some genes/cell types will be amenable. Downregulation of Polθ achieved here by siRNA treatment may alternatively be facilitated by small molecule drugs currently being developed to inhibit repair by Polθ that enables cancer cells to survive therapies that damage DNA.

The reciprocal end-joining pathway identified here supports IHR at frequencies of approximately 0.5% in control cells and 2% in POLQ-depleted cells. Frequencies in this range are anticipated to be useful for gene therapy, especially if corrected cells have an advantage in engraftment or proliferation. IHR has been documented in mouse embryonic stem cells and human B-lymphoblastoid cell lines^18–21^, but at much lower frequencies (<0.01%) which likely reflect targeting of DSBs to only a single homolog in those experiments. End-joining was recently postulated to support IHR that repaired compound heterozygous mutations associated with mouse models of hereditary tyrosinemia type 1 and mucopolysaccharidosis type I^22^. It may be possible to stimulate end-joining to correct mutations found in a variety of mammalian cell types by IHR.

## METHODS

Methods are available in Supplemental Materials.

## ACKNOWLEDGEMENTS

We thank Peter Byers, Hans Ochs and Marshall Horwitz for valuable discussions. This research was supported by grants from the National Institutes of Health to N.M. (R01 CA183967 and R21 CA190675).

## AUTHOR CONTRIBUTIONS

L.D. and N.M. conceived the study and designed experiments. L.D. and K.J.K. carried out experiments. N.M., L.D. and K.J.K. analyzed data. N.M. and L.D. wrote the original draft with critical review and revision by K.J.K.

## REFERENCES

1. Hirschhorn, R. In vivo reversion to normal of inherited mutations in humans. J Med Genet 40, 721–728 (2003).

2. Kane, M.S. et al. Mitotic Intragenic Recombination: A Mechanism of Survival for Several Congenital Disorders of Glycosylation. Am J Hum Genet 98, 339–346 (2016).

3. Lim, Y.H., Fisher, J.M. & Choate, K.A. Revertant mosaicism in genodermatoses. Cell Mol Life Sci 74, 2229–2238 (2017).

4. Rasheed, S., Nelson-Rees, W.A., Toth, E.M., Arnstein, P. & Gardner, M.B. Characterization of a newly derived human sarcoma cell line (HT-1080). Cancer 33, 1027–1033 (1974).

5. Becerikli, M. et al. Growth rate of late passage sarcoma cells is independent of epigenetic events but dependent on the amount of chromosomal aberrations. Exp Cell Res 319, 1724–1731 (2013).

6. Chen, E. et al. RECQL5 suppresses oncogenic JAK2-induced replication stress and genomic instability. Cell Rep 13, 2345–2352 (2015).

7. Deriano, L. & Roth, D.B. Modernizing the nonhomologous end-joining repertoire: alternative and classical NHEJ share the stage. Annu Rev Genet 47, 433–455 (2013).

8. Ceccaldi, R., Rondinelli, B. & D’Andrea, A.D. Repair Pathway Choices and Consequences at the Double-Strand Break. Trends Cell Biol 26, 52–64 (2016).

9. Sfeir, A. & Symington, L.S. Microhomology-Mediated End Joining: A Back-up Survival Mechanism or Dedicated Pathway? Trends Biochem Sci 40, 701–714 (2015).

10. Beagan, K. & McVey, M. Linking DNA polymerase theta structure and function in health and disease. Cell Mol Life Sci 73, 603–615 (2016).

11. Black, S.J., Kashkina, E., Kent, T. & Pomerantz, R.T. DNA Polymerase theta: A Unique Multifunctional End-Joining Machine. Genes (Basel) 7 (2016).

12. Wood, R.D. & Doublie, S. DNA polymerase theta (POLQ), double-strand break repair, and cancer. DNA Repair (Amst) 44, 22–32 (2016).

13. Wyatt, D.W. et al. Essential Roles for Polymerase theta-Mediated End Joining in the Repair of Chromosome Breaks. Mol Cell 63, 662–673 (2016).

14. Schimmel, J., Kool, H., van Schendel, R. & Tijsterman, M. Mutational signatures of non-homologous and polymerase theta-mediated end-joining in embryonic stem cells. EMBO J 36, 3634–3649 (2017).

15. Zelensky, A.N., Schimmel, J., Kool, H., Kanaar, R. & Tijsterman, M. Inactivation of Pol theta and C-NHEJ eliminates off-target integration of exogenous DNA. Nat Commun 8, 66 (2017).

16. Ceccaldi, R. et al. Homologous-recombination-deficient tumours are dependent on Poltheta-mediated repair. Nature 518, 258–262 (2015).

17. Mateos-Gomez, P.A. et al. Mammalian polymerase theta promotes alternative NHEJ and suppresses recombination. Nature 518, 254–257 (2015).

18. Moynahan, M.E. & Jasin, M. Loss of heterozygosity induced by a chromosomal double-strand break. Proc Natl Acad Sci U S A 94, 8988–8993 (1997).

19. Stark, J.M. & Jasin, M. Extensive loss of heterozygosity is suppressed during homologous repair of chromosomal breaks. Mol Cell Biol 23, 733–743 (2003).

20. Neuwirth, E.A., Honma, M. & Grosovsky, A.J. Interchromosomal crossover in human cells is associated with long gene conversion tracts. Mol Cell Biol 27, 5261–5274 (2007).

21. Larocque, J.R. et al. Interhomolog recombination and loss of heterozygosity in wild-type and Bloom syndrome helicase (BLM)-deficient mammalian cells. Proc Natl Acad Sci U S A 108, 11971–11976 (2011).

22. Wang, D. et al. Cas9-mediated allelic exchange repairs compound heterozygous recessive mutations in mice. Nat Biotechnol 36, 839–842 (2018).

23. Davis, L., Zhang, Y. & Maizels, N. Assaying repair at DNA nicks. Meth Enzymol (2018).

24. Davis, L. & Maizels, N. Two distinct pathways support gene correction by single-stranded donors at DNA nicks. Cell Rep 17, 1872–1881 (2016).

25. Mali, P. et al. CAS9 transcriptional activators for target specificity screening and paired nickases for cooperative genome engineering. Nat Biotechnol 31, 833–838 (2013).

