## Supplemental Materials for "Donor-Free Gene Correction by Targeted Interhomolog Recombination"

**Correction of autosomal dominant (“gain of function”) mutation by copy-neutral LOH**

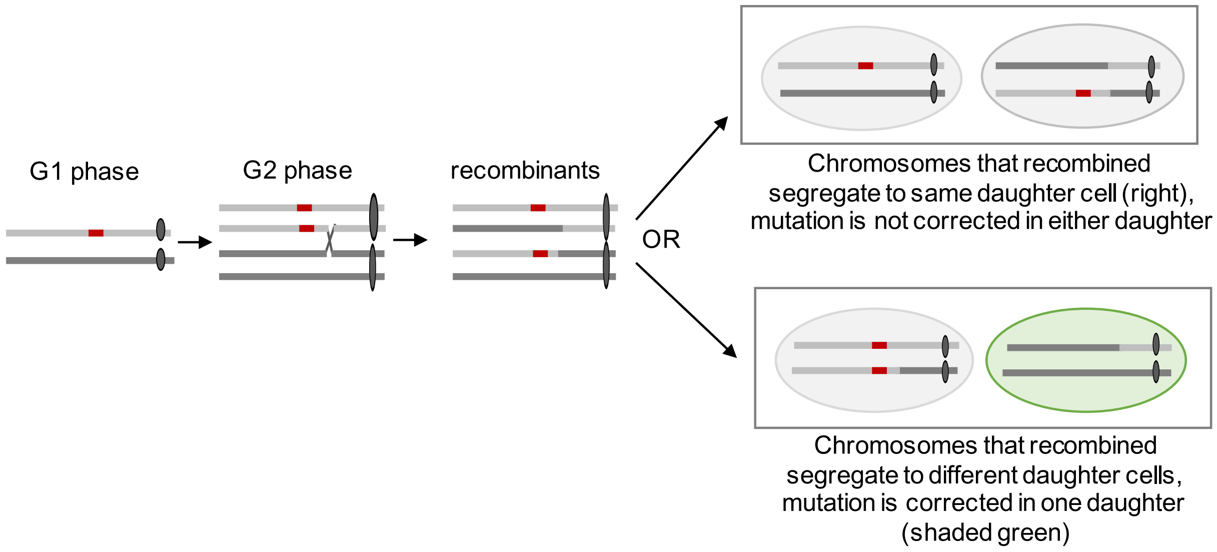

**Supplementary Figure. 1. Correction of an autosomal dominant ("gain of function") mutation by IHR.**

An autosomal dominant mutation (red) is present on one allele. Following replication, crossover recombination (X) at a position between the mutation and the centromere generates one recombinant homolog without a mutation. If that chromosome segregates to the same cell as the chromosome that carried a normal allele in the parental cells, then that segregant will express functional protein (lower right, green shading).

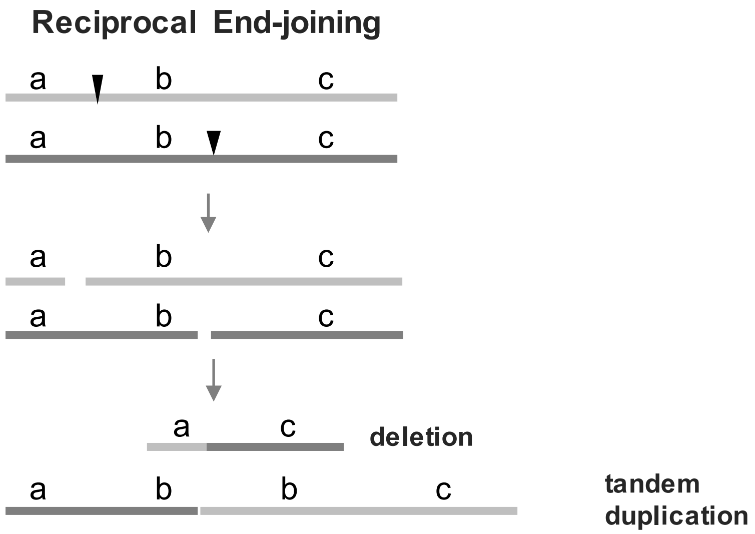

**Supplementary Figure. 2. Reciprocal end-joining at offset DSBs.**
Diagram of predicted recombinants of IHR via reciprocal end-joining at offset DSBs targeted to sites between regions a and b, or b and c. Region b, flanked by the DSBs, will be deleted in one recombinant homolog and exhibit tandem duplication in the other.

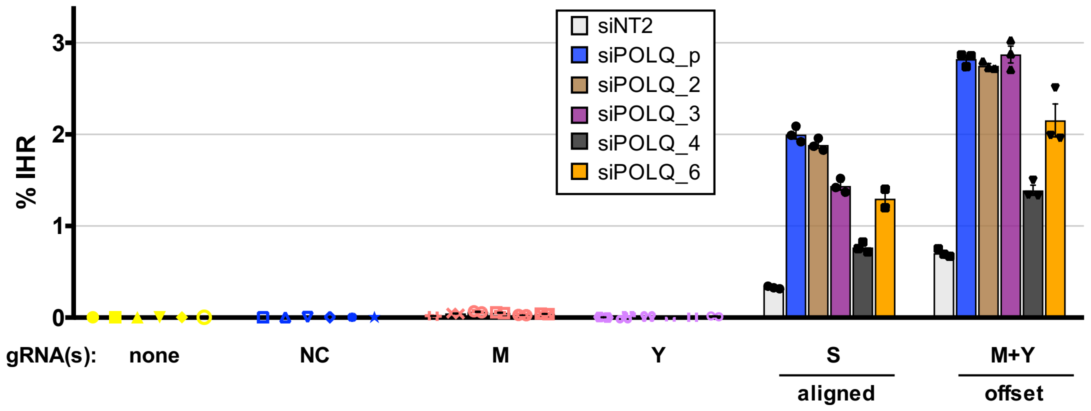

**Supplementary Figure. 3. Effect on IHR of depletion of POLQ by individual and pooled siRNAs.**

IHR frequencies in HT1080-K2 cells treated with siNT2 or with a pool of four siRNAs designed for POLQ depletion (siPOLQ_p, Qiagen catalog # SI02665215, SI00090083, SI00090076, SI00090069), or each of those four siRNAs individually (siPOLQ_6, #SI02665215; siPOLQ_4, #SI00090083; siPOLQ_3, #SI00090076; si-POLQ_2, #SI00090069); and subsequently transduced with RNPs containing no gRNA (-); non-targeting gRNA NC; gRNAs M or Y which cleave on a single allele (**Fig. 1e**); gRNAs M+Y which together generate DSBs offset by 0.6 kb; or S, which cleaves on both alleles (**Fig. 1b**).

### METHODS

#### Data Availability

The datasets generated during and/or analyzed during the current study are available from the corresponding author on reasonable request.

#### Cell culture, staining and flow cytometry

Cells were cultured at 37$^{o}$C, 5% $CO_2$ in DMEM/FBS [Dulbecco-modified Eagle's medium (Hyclone) supplemented with 10% fetal bovine serum (Gemini Bio-Products) and 200 units/ml penicillin, 200 mg/ml streptomycin (Hyclone) and 2 mM L-glutamine (Hyclone)]. For cell storage, approximately 1 x $10^6$ cells were resuspended in 1 ml DMEM containing 10% DMSO and frozen at -80°C.

For analysis of sCD44+ cells frequencies, cells were cultured for 4 days then removed from plates in TrypLE Select (ThermoFisher), fixed by addition of formaldehyde (2% final concentration), and stored at 4°C for up to a week. For staining, fixed cells were pelleted by centrifugation at 1000 rpm for 5 min, the supernatant aspirated, and cells resuspended in 1 ml Dulbecco's Phosphate Buffered Saline (DPBS; HyClone), then pelleted and resuspended in 1ml DPBS containing 1% FBS (DPBSF) at 4°C, then pelleted and resuspended in 300 µl of 1:200 dilution of FITC-conjugated anti-CD44 antibody (clone G44-26; BD Bioscience) in DPBSF. Cells were gently mixed at 4°C for 1 hr, pelleted, and resuspended in 100-300 µl DPBSF for analysis by flow cytometry. For flow cytometry, cells were gated for single cells and CD44-FITC detected with a 488 nm laser on an LSRII flow cytometer (Becton-Dickinson), as previously described[23]. sCD44- control populations were used to gate sCD44+ expression.

To stain cells for subsequent cell sorting, cells were removed from plates, TrypLE Select was inactivated by addition of 5 volumes of fresh DMEM/FBS, and cells were pelleted and washed and stained as above. Single cells were sorted into 100 µl of conditioned DMEM/FBS in 96-well plates using an Aria III cytometer (Becton-Dickinson) with a 488 nm laser.

#### Engineering HT1080 CD44-/- cells to report on IHR

The CD44 gene of the human HT1080 fibrosarcoma cell line[4] was engineered to generate an IHR reporter. HT1080 cells were used as a model because they express functional P53; they propagate with a stable near-diploid karyotype, which avoids complications due to polyploidy; and they have a high cloning efficiency, which facilitates molecular analysis of individual recombinants. The starting population of HT1080 cells (ATCC; Manassas, VA) was confirmed to be near diploid and sCD44+ by flow cytometry of cells stained with DAPI and with anti-CD44 antibody, as described above. This population and its engineered derivatives exhibited a 24 hr doubling time in culture.

To create mutations, DSBs or nicks were targeted by transfection with the indicated gRNA and plasmid constructs expressing Cas9 or Cas9D10A, as previously described[24]. Cells were cultured 4-7 days then single cells sorted into 96 well plates containing 100 μl conditioned medium and expanded for further analysis. Flow cytometry, genotyping and cell storage were carried out as described above.

Mutations in exons 1 and 17 were engineered in 3 steps:

(1) DSBs were targeted to exon 1 by gRNA-CD44-4, single sCD44- cells were sorted and expanded, and exon 1 analyzed by PCR with primers CD44-F2 and R2 and Sanger sequencing. Clone HT1080-G1-C11, in which CD44 exon 1 was mutated on both alleles, was used for subsequent engineering.

(2) The mutation in one allele of exon 1 was corrected by HDR at a nick targeted to exon 1 using gRNA-CD44-D1-2 and supported by SSO donor CD44-1-TOP. Single sCD44+ cells were sorted and expanded, and exon 1 analyzed by PCR with primers CD44-F2 and R2 and Sanger sequencing. Clone HT1080-B7, heterozygous at exon 1 (CD44+/-), was used for subsequent engineering.

(3) A deletion in CD44 exon 17 was engineered by HDR at a nick targeted to this exon with gRNA-CD44T-17-1 supported by SSO donor CD44T-17eng1, which replaced 38 bp of exon 17 with a 17 bp insertion. Single sCD44- clones were sorted and expanded, and exon 17 analyzed by PCR with primers CD44-F4 and R4 and Sanger sequencing. Clone HT1080-K1 (sCD44-) was shown to bear compound heterozygous mutations at CD44 exons 1 and 17. We confirmed that HDR at either exon 1 or exon 17 could restore expression of CD44 in HT1080-K1 cells (sCD44+), using gRNA-CD44-K1B and SSO donor CD44-1-TOP3 (exon 1) and gRNA-CD44T-17-2 and SSO donor CD44T-17-0 (exon 17). HT1080-K1 cells were used to determine IHR frequencies in experiments shown in **Fig. 1d**.

Mutations were engineered in intron 1 of one allele of CD44 to enable analysis of IHR targeted by DSBs on a single homolog and by offset DSBs. Mutagenesis was carried out by HDR at nicks targeted to intron 1 using gRNA W and supported by SSO CD44-intron1-1, designed to cause 3 single base insertions that create recognition sequences for gRNAs Y and Z as well as a diagnostic HindIII site, and to prevent cleavage targeted by gRNA W or X. Following HDR-mediated engineering, single cells were sorted and, expanded; the targeted region of intron 1 amplified with PCR with primers CD44-F16 and CD44-R14, and clones screened for the presence of the diagnostic HindIII site in the modified allele. Mutations was then confirmed by Sanger sequencing. Clone HT1080-K2 was used for analysis of IHR by offset and aligned DSBs in **Fig. 1f** and **Fig. 2 a-c**.

Sequences of the mutant alleles and diagnostic restriction sites are shown in **Supplementary Table 1**.

#### Plasmids

The Cas9 expression construct (pCas9(wt)-T2A-mTagBFP) was previously described[24]. All gRNA expression plasmids were constructed by cloning annealed and extended oligos (Integrated DNA Technologies) into AflII digested gRNA-Cloning Vector[25]. The DNA sequence of the protospacer and PAM for each gRNA are given in **Supplemental Table 2.**

#### IHR assays

IHR was initiated by targeted DSBs generated by transfection of Cas9 and gRNA expression plasmids (**Fig. 1c, d**) or by transduction with Cas9/gRNA RNPs **(Fig. 1f, 2a-c).** For transfections in 24-well plate format, cells were seeded at 5 x $10^4$ cells/well in 0.5 ml DMEM/FBS, incubated overnight, then transfected the following day. Each well received a transfection mix made by combining 150 ng of Cas9-expression plasmid and 75 ng gRNA-expression plasmid (either 75 ng of a single expression construct or 37.5 ng each of two different constructs) in 100 µl Opti-MEM (Thermo Fisher Scientific) reduced serum medium and 1.2 µl Lipofectamine LTX (Thermo Fisher Scientific), and pre-incubated at room temperature for 20 min prior to addition to the well. Cells were cultured 1-2 days, expanded into 6-well plates, further cultured and then collected for analysis at 4 days post-transfection. Transfection efficiencies were reproducibly in the 25-35% range, and IHR frequencies were calculated by dividing the observed frequencies of sCD44+ cells by 0.3. For transductions, Cas9/CRISPR RNPs (crRNA, tracRNA and Cas9; Integrated DNA Technologies, IDT) were used according to the manufacturer's protocol but at amounts corresponding to 25% of the scaled amounts recommended by the manufacturer. Flow cytometry of cells transduced with RNP containing fluorescently labeled tracRNA indicated that transduction frequencies were >95%.

To analyze effects of siRNA depletion on IHR frequencies (**Fig. 2a-c**), cells were seeded in 10 cm plates at 1.8 x $10^6$ cells/plate in 11 ml DMEM/FBS; 2-4 hr later, 1.8 ml of Opti-MEM containing 60 pmol siRNA and 22.5 µl Lipofectamine RNAiMAX (Thermo Fisher Scientific) were added to each plate; and the next day cells were transduced with Cas9/gRNA RNPs. For 24-well plates, 1.5 x $10^5$ siRNA-treated cells in 0.6 ml of DMEM/FBS were added to wells containing 75 µl of Opti-MEM, 4.5 pmol of Cas9 RNP and 1.8 µl of Lipofectamine RNAiMAX. Following 24-36 hr incubation, cells were expanded into 6-well plates. Four days after transfection, cells were collected for analysis. Transfections and transductions were also scaled to 6 cm (siRNA) and 48-well or 6-well plates (RNP), without evident effect on IHR frequencies.

Sample size (n) is indicated in the figure legends and represent a single measurement from each sample. Statistical significance was determined by two-tailed t-test.

**siRNA Depletion**

The siRNAs used were the non-targeting siRNA, siNT2 (ThermoFisher ID #4390847); siRNA targeting BRCA2 (ThermoFisher ID # s2085), validated in physiological HDR assays[24]; siRNA targeting RAD52 (Qiagen catalog # SI03041808, SI03035123, SI03020794, SI02629865); DNA-PKcs (Qiagen catalog # SI02224229, SI02224236, SI02663633); LIG4 (Qiagen catalog # SI03048878, SI03021179, SI00036043, SI00036022) and POLQ (Qiagen catalog # SI02665215, SI00090083, SI00090076, SI00090069). For knockdown of genes with multiple siRNAs shown above, equimolar pools were used. Effects of the pool of four siRNAs designed for POLQ depletion were confirmed by testing each individually (**Supplementary Fig. 3**).

**Single cell expansion, DNA preparation and genotyping**

At 7-10 days after transfection or transduction, single sCD44+ cells were sorted into 100 µl conditioned medium in 96-well plates and cultured for 10-14 days. Cells were then transferred to 24-well plates and cultured an additional 4-6 days. Media was then aspirated and cells were dissociated by adding 105 µl TrypLE™ Select (ThermoFisher) per well. For DNA preparation, 35 µl of this cell suspension was added to 100 µl DMEM to inactivate the TrypLE Select, centrifuged, washed once with PBS, and genomic DNA prepared using DirectPCR Lysis Reagent (Viagen Biotech; Los Angeles, CA).

LOH was analyzed by amplification of exon 1 of CD44 in a 30 µl reaction using primer pair CD44-F2 and CD44-R2, Taq DNA polymerase and ThermoPol buffer (NEB). Restriction digestion was carried out by adding 12 µl of PCR reaction product to 3 µl 1x ThermoPol buffer containing 1.75 U (0.35 µl) Tth111I (NEB; cleaves 5'-GACN↓NNGTC-3') then incubating at 65°C for 90 min. Undigested and digested DNAs were analyzed by 1.3% agarose gel electrophoresis. Similar analyses were performed at exon 17 (CD44-F4 and R4, HindIII or ApoI), rs85074 (rs85074-F and R, BstBI) and rs7950932 (rs7950932-F and R, BsaHI).

Analysis of offset IHR junctions was by amplification of intron 1 using primer pairs CD44-F7 and CD44-R5, CD44-F16 and CD44-R14, or CD44-F17 and CD44-R15. DNA was analyzed by 1.1% agarose gel electrophoresis. Primer pair F15/R12 was used to specifically amplify PCR products spanning the junction of the predicted tandem duplications. All primer sequences are shown in **Supplementary Table 3**.

**Supplementary Table 1. Genomic sequence of relevant regions of CD44 exon1 and intron 1 in the parental HT1080 cell line and its mutant derivatives.**

| cell line | region | sequence |
|---|---|---|
| HT1080 | exon $1^{wt}$ | ATG GAC AAG TTT TGG TGG CAC GCA GCC TGG GGA CTC TGC CTC GTG CCG CTG AGC CTG GCG CAG ATC |
| G1-C11 | exon 1-1 | ATG GAC AAG TTT TGG TGG CAC GCA GCC TGG GGA CTC TGC CTC GTG gCC GCT GAG CCT GGC GCA GAT C |
| | exon 1-2 | ATG GAC AAG TTT TGG TGG CAC GCA GCC TGG GGA CTC TGa CCT CGg TGC CCT GAG CCT GGC GCA GAT C |
| B7 | exon $1^{wt-1}$ | ATG GAC AAG TTT TGG TGG CAC GCA GCC TGG GGA CTC TGt CTC GTG CCG CTG AGC CTG GCG CAG ATC |
| | exon 1-2 | as in G1-C11, above |
| K1 | exon 1 and exon 1-2 | both as in B7, above |
| | exon 17 | AA TGG CTG ATC ATC TTG GCA TCC CTC TTG GCC TTG GCT TTG ATT CTT GCA GTT TGC ATT GCA GTC AAC AGT CGA AGA AG |
| | exon 17-1 | AA gct tga att tag cta gcG ATT CTT GCA GTT TGC ATT GCA GTC AAC AGT CGA AGA AG |
| K2 | exon 1 | as in HT1080-K1, above |
| | exon17 | as in HT1080-K1, above |
| | intron 1-1 | AGACTCTGTTGGATCCAtGGtACCtAGGAGTGGGGGTTAAA |

lower case in red font, insertion

Diagnostic restriction sites (underlined):
Tth111I site (GACNNNGTC) in B7 exon $1^{wt-1}$,
HindIII site (AAGCTT) in K1 exon 17-1;
KpnI site (GGTACC) in K2 intron 1-1.

**Supplementary Table 2. Genomic sequence (+ strand) of gRNA targets.**

| gRNA | location | sequence | plasmid or crRNA |
|---|---|---|---|
| CD44-4 | exon 1 | CCTcgtgccgctgagcctggcgc | plasmid |
| CD44-D1-2 | exon 1-1 | CCTcgtggccgctgagcctggcg | plasmid |
| CD44T-17-1 | exon 17 | accagaatggctgatcatctTGG | plasmid |
| CD44-K1B | exon 1-2 | CCCtgagcctggcgcagatcggt | plasmid |
| CD44T-17-2 | exon 17 | CCAgaatggctgatcatcttggc | plasmid |
| B | intron 1 | ggatgcgcacagtcgttgtcTGG | plasmid |
| F | intron 1 | CCTatgcaactagggtagcctga | plasmid |
| E | intron 2 | CCTgagctacatgagccgttgct | plasmid |
| S | intron 7 | cccctccagagcttaatctaTGG | plasmid and crRNA |
| M | intron 1 | CCAgtgcatcccactcccagat | plasmid and crRNA |
| W | intron 1 | CCAggaccaggagtgggggtta | plasmid |
| X | intron 1 | actctgttggatccaggaccAGG | plasmid |
| Y | intron 1 | CCAtggtacctaggagtgggggg | plasmid and crRNA |
| Z | intron 1 | ctgttggatccatggtacctAGG | plasmid |
| V | intron 1 | CCTgactttcccacttggatgac | plasmid |
| NC | none | IDT catalog # 1072545 | crRNA |

lower case, protospacer
UPPER CASE, PAM (protospacer adjacent motif)

**Supplementary Table 3. Primer and donor oligonucleotide sequences.**

| oligo | region | sequence |
|---|---|---|
| CD44-F2 | exon 1 | 5'-CTTGCTTGGGTGTGTCCTTC-3' |
| CD44-R2 | exon 1 | 5'-CCAAATGGTGCTTCCACAGAC-3' |
| CD44-F4 | exon 17 | 5'-GGCTGTTGGACAAATACCTTCA-3' |
| CD44-R4 | exon 17 | 5'-CTGTCTCTAAAAACCGGGGC-3' |
| CD44-F7 | intron 1 | 5'-ACGTGGCAAGAATAGCAAAGC-3' |
| CD44-R5 | intron 1 | 5'-GGCCAGAATCACACTTGGTG-3' |
| CD44-F16 | intron 1 | 5'-CTGTCCCAGGTATGCACATCA-3' |
| CD44-R14 | intron 1 | 5'-GCAGTCAACGCTGAACCAAC-3' |
| CD44-F17 | intron 1 | 5'-GGCTGCTTTTGGGTGTGAAA-3' |
| CD44-R15 | intron 1 | 5'-TGATGGGCTCCTGGTGAAAG-3' |
| CD44-F15 | intron 1[TD] | 5'-CCACATCCTGGGTTTGTGCT-3' |
| CD44-R12 | intron 1[TD] | 5'-GGTTGGGCCACTTGGTTTTG-3' |
| rs85074-F1 | rs85074 | 5'-GTGACTACAGACCAGCTGATACA-3' |
| rs85074-R1 | " | 5'-GGGCGATCCTTTCATCGTCA-3' |
| rs7950932-F1 | rs7950932 | 5'-AGATGTAGGGCACTGGACTGT-3' |
| rs7950932-R1 | " | 5'-AGAGCCGTACGTGGCTTTTC-3' |
| CD44-1-TOP | exon 1 (silent mutations) | 5'-CGGACACCATGGACAAGTTTTGGTGGCAtGCcGCaTGGGGACTCTGtCTCGTGCCGCTGAGCCTGGCGCAGATCGGTGAGTGCCCGCCGCAGCCTGGGC-3' |
| CD44T-17eng1 | exon 17 (insertion/ deletion) | 5'-CTGAAGCTCACGCATGTCATTTAATTTACTCATACCAGAAgcttgaatttagctagcGATTCTTGCAGTTTGCATTGCAGTCAACAGTCGAAGAAGg-3' |
| CD44-1-TOP3 | exon 1 (wt) | 5'-ACCATGGACAAGTTTTGGTGGCACGCAGCCTGGGGACTCTGCCTCGTGCCGCTGAGCCTGGCGCAGATCGGTGAGTGCCCGCCGCAGCCTGGGCAGCAA-3' |
| CD44-17-0 | exon 17 (wt) | 5'-CTGAAGCTCACGCATGTCATTTAATTTACTCATACCAGAATGGCTGATCATCTTGGCATCCCTCTTGGCCTTGGCTTTGATTCTTGCAGTTTGCATTGCAGTCAACAGTCGAAGAAGG-3' |
| CD44-intron1-1 | intron 1 | 5'-GGGACATATACTTTCTTTTGCCAGAAAGACTCTGTTGGATCCAtGGtACCtAGGAGTGGGGGGTTAAACAGTTCTCCATACCCTCACCTCCAAG-3' |

lowercase, mutations
